# Clarifying the role of higher-level cortices in resolving perceptual ambiguity using Ultra High Field fMRI

**DOI:** 10.1101/2020.05.27.119677

**Authors:** Logan Dowdle, Geoffrey Ghose, Kamil Ugurbil, Essa Yacoub, Luca Vizioli

**Affiliations:** Center for Magnetic Resonance Research, University of Minnesota 2021 6th St SE, Minneapolis, MN 55455

**Keywords:** Top-down, fMRI, Ultra High Field, Cognitive Neuroscience, Vision, Perception, Cortical Network, Task modulations

## Abstract

The brain is organized into distinct, flexible networks. Within these networks, cognitive variables such as attention can modulate sensory representations in accordance with moment-to-moment behavioral requirements. These modulations can be studied by varying task demands; however, the tasks employed are often incongruent with the postulated functions of a sensory system, limiting the characterization of the system in relation to natural behaviors. Here we combine domain-specific task manipulations and ultra-high field fMRI to study the nature of top-down modulations. We exploited faces, a visual category underpinned by a complex cortical network, and instructed participants to perform either a stimulus-relevant/domain-specific or a stimulus-irrelevant task in the scanner. We found that 1. perceptual ambiguity (i.e. difficulty of achieving a stable percept) is encoded in top-down modulations from higher-level cortices; 2. the right inferior-temporal lobe is active under challenging conditions and uniquely encodes trial-by-trial variability in face perception.

## Introduction

Decades of human and animal research has demonstrated that cortical areas are functionally and anatomically linked to form distinct brain networks. Understanding the distinct contributions of individual areas within these networks has been challenging in part because of the rich and reciprocal pattern of connections between brain areas (Alexander et al., 1986; Felleman and Van Essen, 1991; Moeller et al., 2008). Discrimination is further confounded by the low latencies with which signals travel between areas (Laughlin and Sejnowski, 2003; Wang et al., 2008), substantial overlaps in the functional sensitivities of neurons of different areas (Arcaro and Livingstone, 2017; Haxby et al., 2001; Vogels and Orban, 1994), and the flexibility of connections necessary to maintain function across a range of tasks and environments (Bassett et al., 2011; Gonzalez-Castillo and Bandettini, 2018; Kabbara et al., 2017). A critical component of this flexibility in sensory systems is top-down control, in which cognitive variables such as attention, value, and memory can alter sensory representations in accordance with explicit or specific behavioral demands. Yet this flexibility also is a major challenge to experimental studies, because it implies that the nature of signal processing both within and between brain areas of a network is not fixed and very much depends on the behavioral context and goals of the subject.

While many humans and animal studies have demonstrated the ability of task manipulations to modulate sensory evoked responses, in most cases these studies have employed tasks that are not specific to the domain of the stimulus that was presented. For example, numerous electrophysiological and imaging studies suggest that attending to a particular location in visual space increases the responses of neurons whose receptive fields lie within the attended location irrespective of the selectivities of those receptive fields (Cohen and Maunsell, 2011; Liu et al., 2015; Martinez-Trujillo and Treue, 2004). Similarly, perceptual tasks involving working memory often result in a broad distribution of enhanced signals in sensory areas (Druzgal and D’Esposito, 2003, 2001; Kay and Yeatman, 2017; Leung and Alain, 2011; Pessoa et al., 2002). By contrast, top-down modulation from domain-specific tasks can be far more targeted, selectively enhancing only those neurons whose receptive field responses are consistent with behavioral demands (De Martino et al., 2015; Zhang and Kay, 2020). Accordingly, the use of appropriate domain specific tasks is likely to highlight those neurons with relevant selectivities and reveal how they contribute to perception. For example, contrasting the activity elicited by identical face stimuli in an N-back and a fixation task may reveal working memory related modulations that are unspecific to face processing and therefore fail to elucidate the changes related to the specialized processing of faces. Indeed, a N-back face task could be performed without explicit face recognition or perception; additionally, one could imagine that explicit vs. implicit face detection may be underpinned by different neural computations, and therefore, implementing a task with no explicit requirement may represent a confound or lead to a different set of results.

Establishing how different areas within a network contribute to perception requires sampling the entire network and then identifying the particular areas that can best explain task performance. However, technical limitations have precluded this approach. Invasive electrophysiological studies are necessarily highly targeted and unable to sample across an entire brain network. In addition, these techniques are frequently performed in non-human primates, with tasks that fail to capture the cognitive sophistication available to humans. Moreover, even whole brain studies, such as those relying upon functional magnetic resonance imaging, are often highly biased in their sampling because of the difficulty of imaging particular areas. For example, a number of high-level areas – crucial for human cognition – are located in cortical loci that are traditionally difficult to image (e.g. due to low SNR and their proximity to air cavities (Chen et al., 2003; Devlin et al., 2000)). These limitations also constrain our ability to establish appropriate animal models for invasive measurements and manipulations because of the possibility that human areas homologous to those in animal studies are difficult to sample with conventional gradient echo fMRI methods (Rajimehr et al., 2009). Signal to noise issues have also limited the ability of fMRI to establish trial-to-trial behavioral covariance, which is a critical requirement for understanding the neural basis of perception (Parker and Newsome, 1998) and has been extensively employed by electrophysiology studies (Desimone et al., 1984; Hubel and Wiesel, 1959; Martinez-Trujillo and Treue, 2004; Perrett et al., 1982). This is a particularly important consideration because, by definition, all areas within a network are co-activated, but, given the richness of connectivity within the network, that co-activation does not necessarily imply perceptual relevance.

While the difficulties surrounding magnetic field inhomogeneity and transmit uniformity can be more detrimental at higher fields, there are substantial SNR gains available with ultra-high field magnets (UHF i.e. 7T and above). In common practice, these SNR gains are generally traded for higher resolutions (e.g. submillimeter). In this paper we instead select a relatively lower resolution (i.e. 1.6 mm^3^) and leverage the improved SNR available in UHF fMRI to study the entire ensemble of brain areas associated with a behaviorally important high-level perceptual task, namely the detection of faces in poor visibility. We choose faces as these are a well-studied, cross species visual category that is highly meaningful and therefore ideal for investigating high-level, top-down neural modulations. The human face processing network has however proved challenging to study (Kanwisher et al., 1997; Kanwisher and Yovel, 2006; McCarthy et al., 1997) (see Grill-Spector & Weiner 2014 for review), because many of its areas, due to their proximity to air cavities, are heavily affected by the aforementioned fMRI signal difficulties. In order to combat susceptibility related dropout we leverage the improved acceleration available at high fields to increase our imaging resolution to 1.6mm relative to the 2mm gold-standard for 3T studies. Combined with manual shimming, this yields substantial gains in SNR, while reducing drop out (Farzaneh et al., 1990; Olman et al., 2009; Young et al., 1988) in key human face processing areas. In the current work, we varied the visibility of face stimuli by modulating their phase coherence (Figure 1) and instructed the subjects to perform 2 tasks in the scanner: a domain-specific, stimulus relevant task involving perceptual judgment of the visual stimuli (i.e. face detection); and, stimulus irrelevant fixation task. By leveraging signal gains derived from our imaging approach, we then contrast these responses to isolate top-down modulations specific to the face domain and link subjective perception to brain activity.

**Figure 1.**
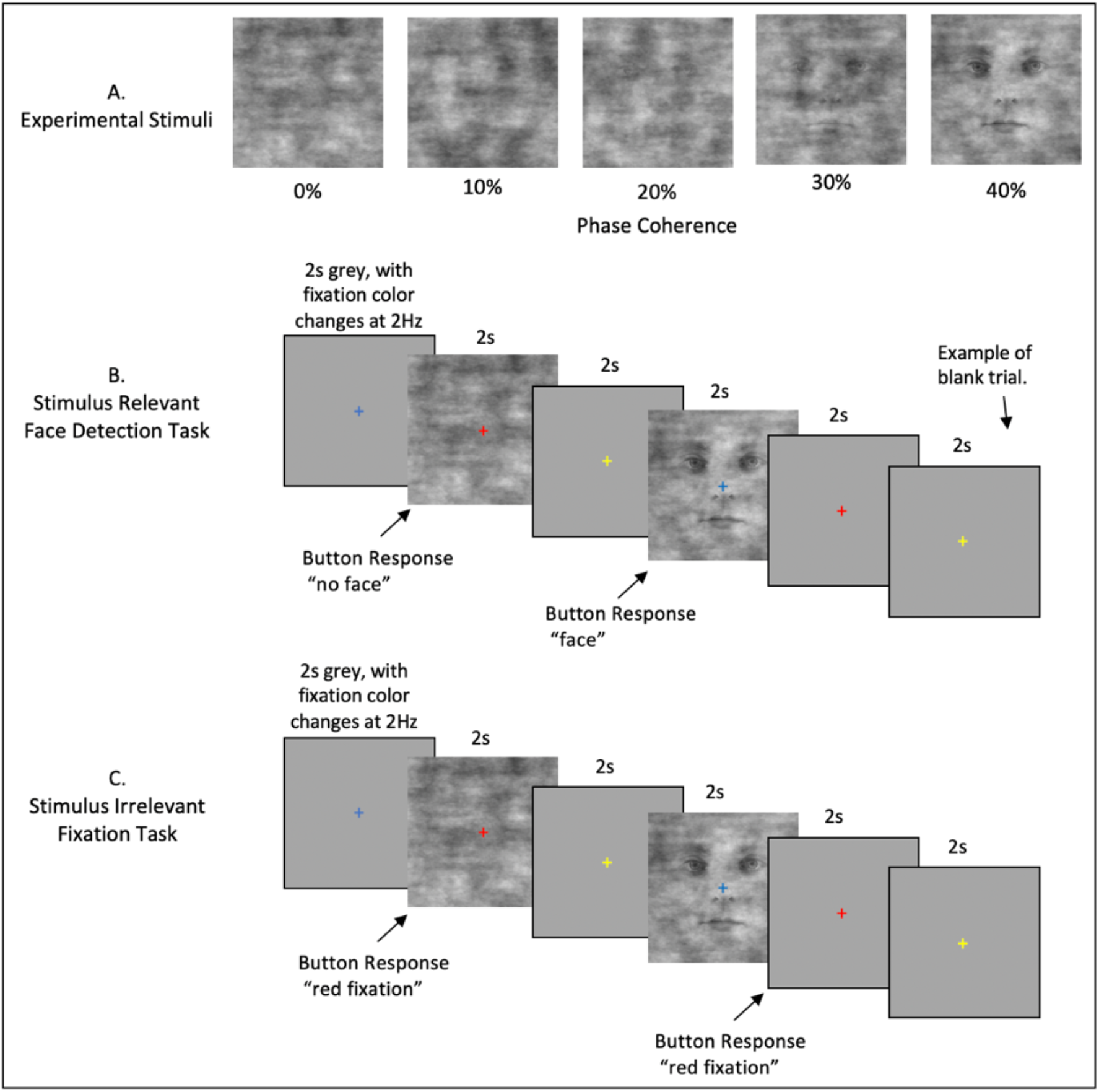
Event Related Stimuli and Tasks. **A)** Example stimuli associated with the 5 phase coherence levels used. **B)** The Stimulus Relevant Face detection task. Stimuli appeared for 2 seconds, with an inter-trial interval of 2 seconds. The fixation cross changed color with a frequency of 2Hz and was visible throughout all experimental procedures. Participants indicated “face” or “no face” using an MRI response hand pad. Blank trials were randomly inserted during each fMRI run. **C)** The Stimulus Irrelevant Fixation task. Timing is identical to B. Participants indicated when the fixation cross changed to the color red.

## Materials and Methods

### Participants

10 (5 females) healthy right-handed subjects (age range: 18-31) participated in the study. Of these, 1 participant was re-scanned due to excessive motion during scanning. All subjects had normal, or corrected vision and provided written informed consent. The local IRB at the University of Minnesota approved the experiments.

### Stimuli and procedure

The experimental procedure consisted of a standard block design face localizer and a fast event-related face paradigm. For both experiments, the stimuli were centered on background of average luminance (25.4 cd/m2, 23.5-30.1). Stimuli were presented on a Cambridge Research Systems BOLDscreen 32 LCD monitor positioned at the head of the 7T scanner bed (resolution 1920, 1080 at 120 Hz; viewing distance ∼89.5 cm) using a Mac Pro computer. Stimulus presentation was controlled using Psychophysics Toolbox (3.0.15) based code. Participants viewed the images though a mirror placed in the head coil. Behavioral responses were recorded using Cambridge Research Systems button box and Psychophysics Toolbox.

### Face localizer

#### Stimuli

All visual stimuli used for the face localizer consisted of grayscale photographs depicting 20 different faces (10 identities × 2 genders) and objects (both taken from Stigliani et al., 2015) and textures of noise.

The experiment consisted of the presentation of grayscale stimuli drawn from different stimulus categories. There were 11 categories, grouped into 3 stimulus domains: faces (adult, child), no face objects (including characters (word, number), body parts (body, limb), places (corridor, house), and objects (car, instrument)) and phase scrambled faces, computed by randomly scrambling the phase coherence of the face stimuli. Each stimulus was presented on a scrambled background (different backgrounds for different stimuli) and occupied a square region with dimensions 10° × 10°. Noise texture stimuli were created by randomly scrambling the phase of the face images (i.e. 0% phase coherence).

The stimuli were equated in terms of luminance, contrast and spatial frequency content by taking the Fourier spectra across stimuli and ensuring that the rotational average amplitudes for a given spatial frequency were equated across images while preserving the amplitude distribution across orientations (Willenbockel et al., 2010). The root mean square contrast (i.e. the standard deviation of pixel intensity) was also kept constant across stimuli.

#### Visual presentation paradigm

Face localizer runs involved presentation of blocks of faces, objects and noise textures. Each run began with presentation of a black fixation cross displayed on a grey background for 12 sec and consisted of 9 randomly presented blocks of images. Each block (3 blocks/category; separated by a 12 sec fixation) involved presentation of 10 different stimuli randomly presented for 800 ms, separated by a 400 ms interstimulus interval (ISI). To ensure that participants’ attention was maintained throughout the localizer we implemented a 1-back task, where subjects were instructed to respond to the repetition of 2 identical stimuli (representing about 10% of the trials), by pressing a button on a response pad held in their right hand. All participants completed two runs of the block design face localizer, where each block occurred 3 times within a run, for a total run duration of 228 seconds.

### Event related experiment

#### Stimuli

We used grayscale images of faces (20 male and 20 female), presenting neutral expressions. We manipulated the phase coherence of each face, from 0% to 40% in steps of 10%, resulting in 200 images (5 visual conditions x 20 identities x 2 genders). We equated the amplitude spectrum across all images. Stimuli approximately subtended 9° of visual angle. Faces were cropped to remove external features by centering an elliptical window with uniform gray background to the original images. The y diameter of the ellipse spanned the full vertical extent of the face stimuli and the x diameter spanned 80% of the horizontal extent. Before applying the elliptical window to all face images, we smoothed the edge of the ellipse by convolving with an average filter (constructed using the “fspecial” function with “average” option in MATLAB – see Figure 1A). This procedure was implemented to prevent participants from performing edge detection, rather than the face detection task, by reacting to the easily identifiable presence of hard edges in the face images.

Like for the localizer experiment, here too we equated amplitude spectrum across the whole stimulus set. We controlled the Fourier spectra across stimuli, ensuring that the rotational average amplitudes for a given spatial frequency were equated across images while preserving the amplitude distribution across orientations (Willenbockel et al., 2010). The root mean square contrast (i.e. the standard deviation of pixel intensity) was also kept constant across stimuli.

#### Tasks

In the scanner, participants were instructed to maintain fixation on a central cross throughout the run and to perform one of 2 tasks: one domain-specific face detection task, involving perceptual judgment of the visual stimuli; and a second, difficult, non-specific attention fixation task that required responding to a specific color change (i.e. red) of the fixation cross.

In the former, participants were instructed to respond as quickly as possible by pressing one of 2 buttons on their button box to indicate whether they perceived a face. Subjects’ instructions were carefully delivered to indicate that there were no correct answers and that we were instead interested in the subjects’ perception only (Figure 1B).

The latter was designed to isolate bottom-up stimulus-driven responses. In order to ensure this, we piloted the fixation task to maximize task difficulty and direct the attention away from the face stimuli on 4 participants that were not included in the experimental subject pool. Based on participants’ feedback, the fixation tasks entailed pressing a button every time the fixation turned red. Every 500 ms, the fixation changed to one of five colors — specifically red, blue, green, yellow and cyan — in a pseudorandom fashion, avoiding consecutive presentations of the same color (Figure 1C). The color change occurred out of sync with the stimulus presentation onset and frequency of button presses was kept constant across tasks. Visual stimuli were identical across tasks to ensure that any differences observed were related solely to top-down processes. Tasks were blocked by run and counterbalanced across participants.

#### Visual presentation paradigm

We acquired 3 runs per task. Each run lasted approximately 3 mins and 22 secs and began and ended with a 12-second fixation period. Within each run, we showed 40 images (5 phase coherence levels x 4 identities x 2 genders) presented for 2000 ms, with a 2000 ms interstimulus interval (ISI). Importantly, we introduced 10% blank trials (i.e. 4000 ms of fixation period) randomly interspersed amongst the 40 images, effectively jittering the ISI. Stimulus presentation was pseudorandomized across runs, with the only constraint being the non-occurrence of 2 consecutive presentations of the same phase coherence level. Behavioral metrics, including reaction time and responses to face stimuli indicating participants’ perceptual judgments (i.e. face or no face) were generated per subject and then averaged.

#### Ambiguity Calculation

For the face detection task, we calculated the average face response percentage to each phase condition, averaged across runs. These values vary from 0 to 100%, representing consistent non-face to consistent face response respectively. Within each subject, this produces a sigmoidal shaped curve. We mathematically define the ambiguity score, on a subject by subject basis, as the inverse of the absolute distance from the inflection point on the sigmoid that fit this curve This is shown in Equation 1:

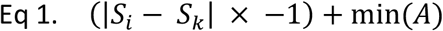

where S_i_ is the *i*th point of the faceness sigmoid, S_k_ is the theoretical midpoint of the faceness sigmoid and min(A) is the minimum value of the ambiguity function. While conceptually similar to defining a perceptual threshold, the goal of this analytical approach is to characterize the instability of the percept in a threshold-free, data-driven manner for all visual conditions, rather than determining the precise level of phase coherence information required to achieve a reliable face percept.

### MR Imaging Acquisition and Processing

All functional MRI data were collected with a 7T Siemens Magnetom System with a 1 by 32-channel NOVA head coil. T2*-weighted images were collected using sequence parameters (TR 1s, Multiband 5, GRAPPA 2, 7/8ths Partial Fourier, 1.6mm isotropic, TE 22ms, Flip Angle 52°, Bandwidth 1923Hz, 85 slices, FOV 208 x 208 mm) matched to the Human Connectome 7T protocol (Thanh Vu et al., 2017).

A number of steps were taken to maximize signal quality of the present study. First, we manually shimmed the B0 field to maximize homogeneity over ventral and anterior-temporal regions. Moreover, we selected our flip angle based on the offset between the selected flip angle and the subject specific flip angle, using the results from a 3dAFI (Actual Flip Angle Image) sequence. In this way we optimized flip angles across the brain to maximize SNR in the ventral and anterior-temporal regions. Finally, by taking advantage of the greater ability to accelerate image acquisition at 7T, we were able to obtain higher resolution images and relative reductions in magnetic susceptibility dropout in these regions (Farzaneh et al., 1990; Olman et al., 2009; Young et al., 1988).

T1-weighted anatomical images were obtained using an MPRAGE sequence (192 slices; TR, 1900 ms; FOV, 256 x 256 mm; flip angle, 9°; TE, 2.52 ms; spatial resolution, .8 mm isotropic voxels) which were collected with a 3T Siemens Magnetom Prisma^fit^ system. Anatomical images were used for visualization purposes only.

#### Functional Image Processing

Dicom files were converted using dcm2niix (Li et al., 2016). Subsequence functional image processing was performed in AFNI version 19.2.10 (Cox, 1996). Conventional processing steps were used, including despiking, slice timing correction, distortion and motion correction, and alignment to each participant’s anatomical image. With each task type, the target for time series and anatomical alignment was the Single Band Reference (SBRef) image which is acquired to calibrate coil sensitively profiles prior to the multiband acquisition and has no slice acceleration or T_1_-saturation, yielding high contrast (Smith et al., 2013). In order to improve localizer and task alignment, an additional nonlinear transform was computed between the task SBRef and localizer SBRef. All spatial transforms were concatenated and applied in a single step to reduce image blurring, and functional images were inspected to confirm successful registration to anatomical targets.

#### Localizer Task Analyses

For the functional localizer, the data was then smoothed with a Gaussian kernel with FWHM of 2 voxels (3.2mm), and each run scaled to have mean 100. For regressors of no interest, the 6 estimated rigid-body motion parameters and polynomials up to order 3 (including linear trend) were added to the design matrix.

#### Functional ROI Definition

Using 3dDeconvolve, the data were passed through a GLM in order to determine the response to faces, objects and noise. Each event was modeled as a 12s box car, convolved with AFNI’s SPMG1 HRF estimation. Rather than select face patches by comparing face activation to the average of objects and noise, we instead constrained the statistical map in the following ways: 1) Betas to faces were positive, 2) the T-stat used for thresholding was the minimum positive T-stat from faces > objects or faces > noise. In other words, we selected voxels that were significant for faces greater than objects, in conjunction with faces greater than scrambled. Regions of interest (ROIs) were derived in volume space, and all consisted of contiguous clusters made up of 19 or more voxels. Statistical thresholding was adjusted to obtain consistency between subjects, no voxels above p<0.05 were considered. To aid in ROI definition, we simultaneously viewed the surface representation of the statistical map using SUMA (Saad and Reynolds, 2012) with FreeSurfer (Dale et al., 1999; Fischl et al., 2004, 2002, 2001) defined cortical surfaces and atlas labels. In addition, to supplement the statistical parametric maps, we used finite impulse response (FIR) deconvolution, with 20 bins, to estimate HRFs in response to each localizer condition (face, scrambled, object). These HRFs were also used to inspect data quality for each ROI.

#### Face Detection and Fixation Color Tasks Analyses

For the Face/Fix detection tasks, the data were only scaled following initial processing, no smoothing was used. As we were interested in BOLD responses across the whole brain, and we know that hemodynamic response functions differ across cortical regions (Handwerker et al., 2004; Lewis et al., 2018; Taylor et al., 2018), we performed a GLM based finite impulse response (FIR) deconvolution analyses, which does not make assumptions regarding the shape of the HRF. This was done using AFNI’s TENTZero method, estimating responses with 1s bins out to 15s post stimulus. Deconvolution was performed separately for each task (face detection and fixation) and phase coherence condition, generating 5 FIR curves for the fixation task and 5 for the face detection task. For each task and independently per subject, ROI, run and condition, we averaged all FIR curves across all voxels. To avoid the contribution of extreme voxels, we trimmed 5% of the values falling at each extreme of the distribution tail to compute the 10% trimmed mean. We then extracted the 4 timepoints corresponding to 4, 5, 6 and 7 seconds after stimulus onset, i.e. those with the largest amplitude within a time window spanning from 2 to 10 TRs after stimulus onset. The 12 extracted amplitudes (4 timepoints per run) were then averaged within experimental tasks to obtain one percent signal change value per subject, ROI, task and condition.

### Statistical Analyses

To test for statistically significant differences between conditions (i.e. phase coherence levels) and tasks, we carried out the following statistical tests.

#### Task and Condition Analyses

Independently per ROI, we performed a 2 (tasks) by 5 (phase coherence levels) repeated measures ANOVA with the mean BOLD response as a dependent variable. To control for family wise error rate (FWER), we implemented the following multiple comparison correction procedure: we began by centering the data on the group mean for each condition and task. This procedure effectively put the data under the ideal H0 hypothesis of no difference between the means. We then sampled with replacement the subjects and performed the same 2 x 3 repeated measures ANOVA and stored all F values for the relevant main effects and interactions. We repeated this procedure 10,000 times and selected the 95% largest F values. We used these F values as our new thresholds and considered statistical significance only when p values for the original ANOVA were smaller than .05 *and* the connected F values were larger than their centered counterpart (e.g. (Wilcox, 2005))

When appropriate (i.e. for comparisons entailing more than 2 factors), we further performed post-hoc paired sample t-tests on significant (as defined above) main effects and interactions. The same FWER control procedure described above for the ANOVA test was implemented to account for multiple comparisons.

Additionally, we performed power analyses to determine effect size and the sample size required to achieve adequate power (see results). Specifically, we computed Hedges’ g (Freeman et al., 1986; Hedges, 1981). This choice was motivated by the fact that, unlike Cohen’s d, which, especially for small samples (i.e. n < 20), tends to provide positively biased estimators of population effect sizes (Freeman et al., 1986; Lakens, 2013), Hedges’ g tends to be unbiased (Cumming, 2012).

#### Functional connectivity analyses

To determine the extent to which task demands modulate functional connectivity amongst face ROIs, we carried out the following analysis. Independently per subject, task and ROI, we averaged (mean) all FIR response curves amongst voxels and concatenated the time courses elicited by each condition to produce a single time course. The concatenation was done to maximize statistical power, as the resulting time course contained 80 timepoints (i.e. 16 FIR timepoints x 5 conditions). For each subject and task, we then computed Pearson’s correlation coefficient between the concatenated time courses of all possible pairs of ROIs. This procedure lead to the formulation of 2 connectivity matrices (12 ROIs x 12 ROIs) per subject (1 per task) summarizing the similarity of BOLD responses between all pairs of ROIs. To convert the skewed sampling distribution of Pearson’s r into a normal distribution we computed Fisher’s z transformation (Fisher, 1915). We therefore proceeded to carry out paired sample t-tests between the connectivity estimated obtained for the 2 tasks. To control for FWER, we implemented the same bootstrap procedure on centered data described in the previous paragraphs (see paragraph 4.6.1). For display purposes only, after computing the mean between the Fisher z-normalized connectivity matrices, we computed the inverse of such transformation on the group average connectivity matrices to convert these scores back into meaningful and interpretable Pearson’s r.

To better visualize the results of our functional connectivity analysis, we further performed classic multidimensional scaling (MDS - using the function “cmdscale” in MATLAB) on the participants average dissimilarity matrix (i.e. 1-Pearson’s r). MDS was performed independently per task. For ease of visual comparison, the MDS arrangements of the 2 tasks were aligned by means of linear transformations (including translation, reflection, orthogonal rotation, and scaling) using Procrustes rotations (this was implemented with the “procrustes” function in MATLAB) with the sum of squared errors as stress metric.

MDS is a useful data driven approach to visualize the data projected into a new space whose dimensions are the first (in this case 2) eigenvectors (i.e. those explaining most of the variance in the data) without any prior hypotheses. MDS is therefore a dimension reduction technique that highlights the dominant features in the data. Results are shown in Figure 4; distance between the points indicate dissimilarity of responses.

**Figure 4.**
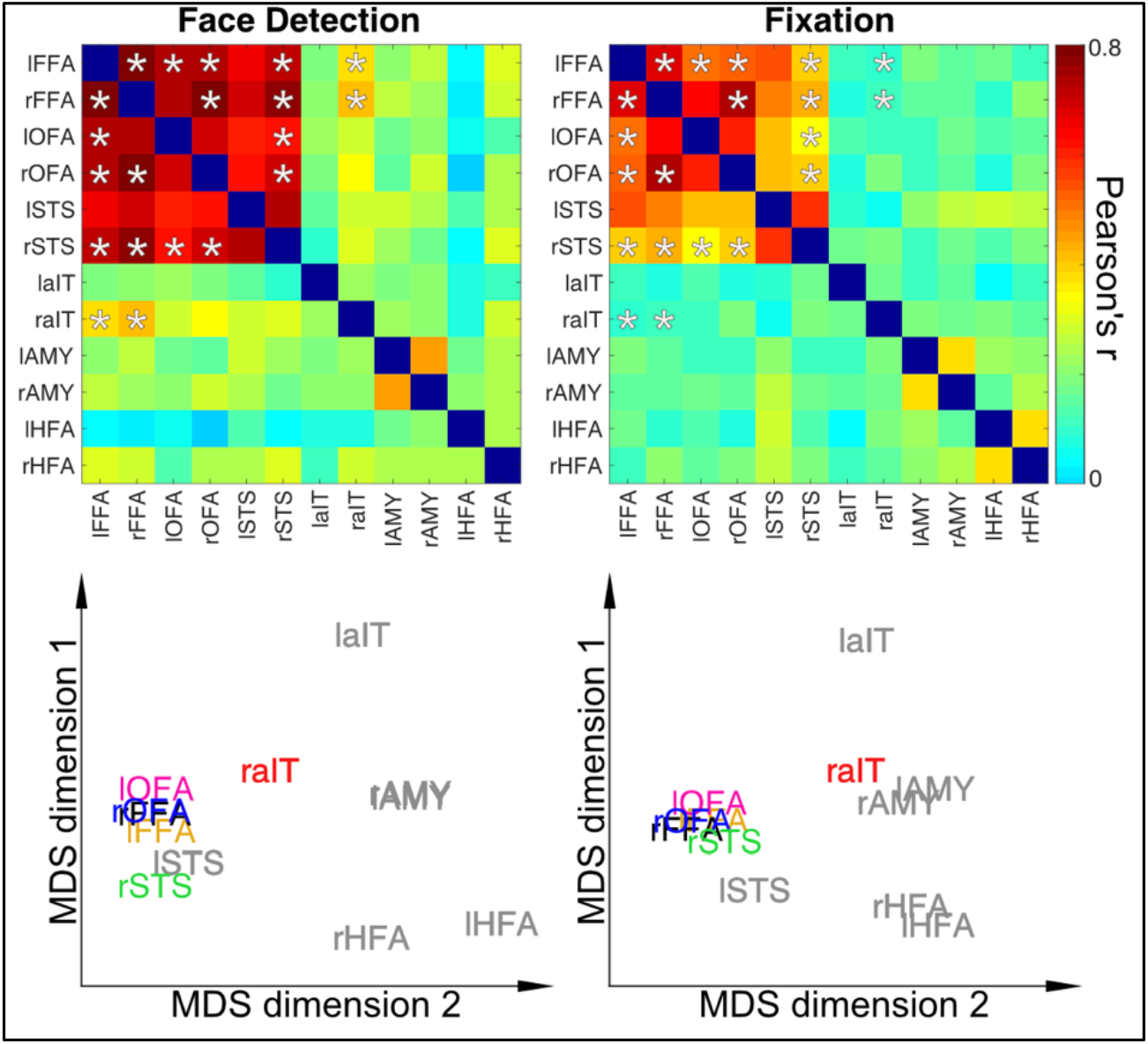
Dynamic Reconfiguration of Face Network as a function of Task **Top Row**: Connectivity Matrices for Stimulus Relevant Face Detection Task (Left) and Stimulus Irrelevant Fixation Task (Right). Asterisks indicate correlation coefficients that significantly (p<0.05) difference between tasks and are identical between matrices for visualization purposes. **Bottom Row**. Classic Multidimensional Scaling for connectivity matrices highlights the higher proximity of the rAIT to the core face areas as a function of increased connectivity during face detection relative to the fixation task. ROIs in grey text indicate those regions with no significant connectivity modulations across tasks.

#### Brain-behavior correlation

Next for each ROI, we wanted to assess the relationship between behavioral responses and top-down BOLD modulations. To this end, for each subject and ROI, we computed Pearson’s correlation coefficients amongst the ambiguity scores at each phase coherence level and the task difference in average BOLD amplitudes elicited by each condition. We then performed Fisher z transform (see paragraph above) on Pearson’s r and carried out one sample t-test for each ROI to determine whether the average group correlation was significantly larger than 0. To control the family wise error rate (FWER), we implemented the same bootstrap procedure on centered data described in the previous paragraphs (see paragraph 4.6.1). For display purposes only, after computing the mean between the Fisher z-normalized correlation scores, we computed the inverse of such transformation on the group average to convert these scores back into meaningful and interpretable Pearson’s r.

#### Perception-Based Analyses

We performed additional analyses to assess BOLD amplitude modulations as a function of percept. Due to the grouping of responses, we considered only the most ambiguous condition (i.e. 20% phase coherence, see results). We allocated each trial to one of 2 new conditions: “*face percept*” and “*no face percept*” on the basis of each participants reported percept. We then repeated the analysis described in section 2.5.4 to estimate BOLD amplitudes of these 2 conditions.

## Results

Through this work, the prefixes “l” and “r” preceding the name of a ROI will refer to its hemispheric lateralization.

### Localizer Task

Using the separate face-localizer task, we identified a total of twelve ROIs for all participants. This included typical regions, such as the fusiform face area (FFA, MNI Centers of Mass [L: -41 -52 -19, R: 40 - 54 -17]), occipital face area (OFA [L: -44 -83 -13, R:43 -76 -11]), posterior superior temporal sulcus (STS [L: -48 -68 10, R: 50 -57 18]) as well as the amygdalae [L: -21 -7 -15, R: 20 -6 -15]. In addition, we identified ROIs for more difficult regions, including an ROI proximal to the perirhinal cortex, which we are referring to more generally as the anterior inferior temporal (AIT [L: -34 -9 -34, R:32 -6 -39]) cortex as well as an ROI in H-shaped sulcus, which we refer to as the HFA (H-shaped sulcus Face Area, [L: -32 33 -15, R: 30 33 -14]). ROIs identification and all subsequent analyses were carried out in native subject space, but to compare their locations across participants and to relate these locations to previous studies, we converted and report their coordinates in MNI space. All cortical regions and their overlap (in the specific slices selected) across participants are shown in Figure 2A.

**Figure 2.**
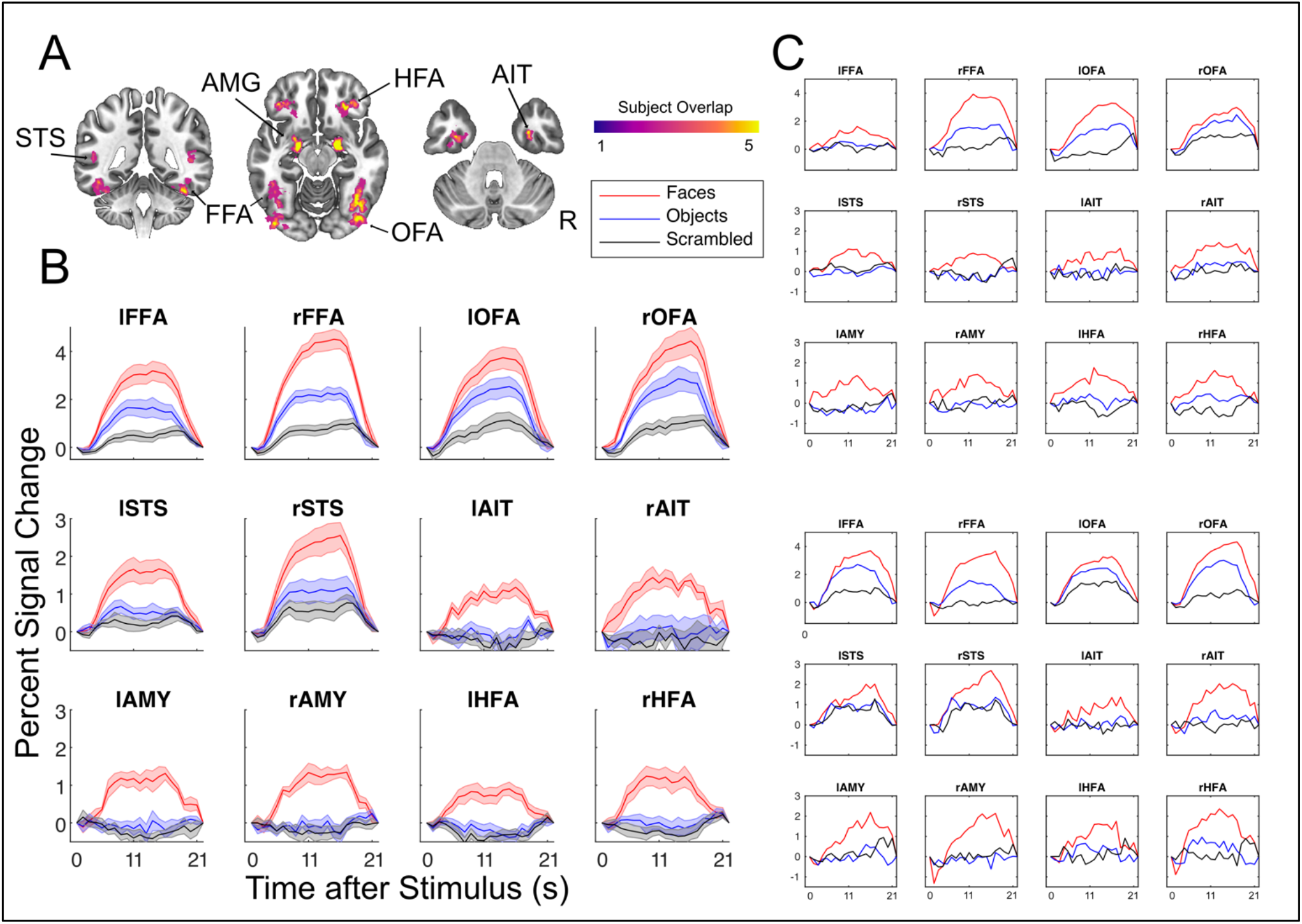
Results from the face preferential localizer. **A)** All subject’s ROIs combined in MNI template space, highlighting the consistent overlap between subjects and between hemispheres. Labels indicate ROI names, right is right. **B)** Average BOLD responses across all subjects, in units of percent signal change, for each ROI in response to faces, objects and scrambled images. Standard Error of the Mean shown with shading. **C)** Example results from two single subjects showcasing reliability even within a single subject’s ROIs.

Figure 2B shows ROIs average BOLD time courses, constructed from FIR models. All regions, including areas associated with low SNR (AIT, HFA), yield plausible hemodynamic responses. Furthermore, these responses remain HRF-like even at the single subject level (Figure 2C), indicating high data quality.

### Behavioral Responses and Ambiguity Scores

The right panel in Figure 3A portrays the face detection group average reaction times. Subjects responded with an average RT of 760 +/- 120 ms. Reaction times indicate that, in reporting their percepts, participants were slowest at the lower end of the phase coherence spectrum (i.e. 0% and 10%), becoming increasingly faster as a function of phase coherence.

**Figure 3.**
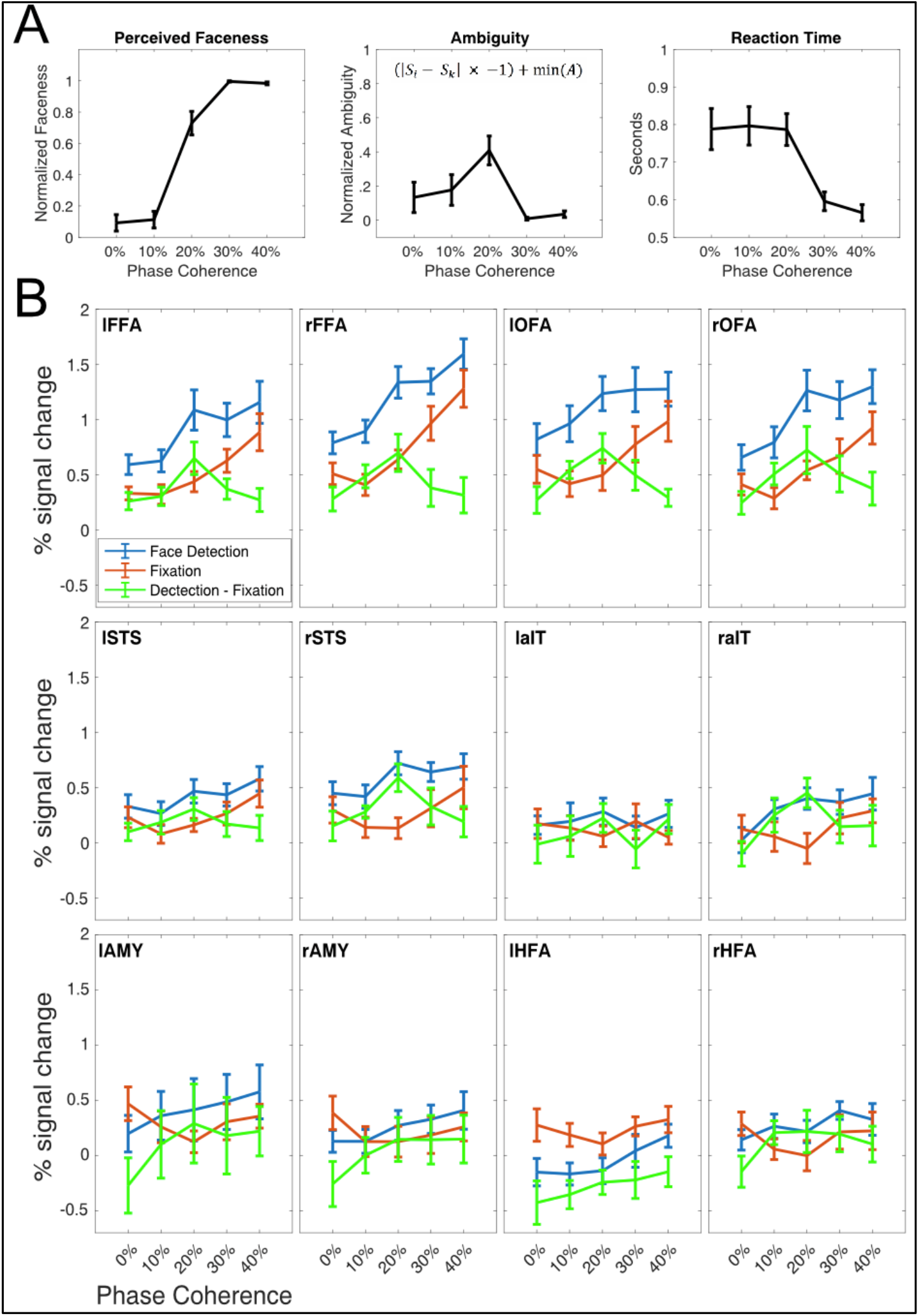
**A)** Behavioral Responses to Stimulus Relevant Task. **Left panel:** The group average of perceived “faceness”, where 0 represents consistently reporting no face, and 1 represents consistently reporting the presence of a face. **Central Panel:** The group averaged ambiguity curve, showing that stimuli at 20% were most ambiguous, i.e., were most inconsistently categorized. Ambiguity was computed according to the equation shown in the middle panel, where S_i_ is the *i*th point of the faceness sigmoid, S_k_ is the theoretical midpoint of the faceness sigmoid and min(A) is the minimum value of the ambiguity function. **Right Panel:** Group averaged reaction times during the face detection task. For all panels, error bars represent standard errors across subjects. **B)** Percent Signal change during event related tasks in all ROIs. In blue, the BOLD responses to the stimulus relevant, domain specific task, in red to the stimulus irrelevant task. For the majority of ROIs, the stimulus relevant BOLD responses are larger across all phase levels relative to the stimulus irrelevant task. The green curve represents the differences between tasks.

Figure 3A, left panel, shows the average perceived “faceness”, that is the proportion of the time that participants reported seeing a face. By calculating the inverse of the absolute distance from the inflection point (see section 4.4.4.) of the faceness sigmoid, we derive the ambiguity scores (Figure 3A, middle panel), which peaked at 20% phase coherence. Reaction times were shortest for the highest phase coherence (Figure 3A, right panel).

### Face and Fixation fMRI analyses

#### ANOVA Results

The subjects mean percent signal change in response to each condition, for each ROI, is shown in Figure 3B.

The main effect of the task was significant (p <0.05) in the lFFA (F_1,9_ = 20.559), rFFA (F_1,9_ = 13.131), lOFA (F_1,9_ = 48.595), rOFA (F_1,9_ = 14.304) and the rSTS (F_1,9_ = 9.182), with face detection driving larger BOLD responses relative to the fixation task.

There was a significant (p <0.05) main effect of the condition (phase coherence level) in the lFFA (F_4,36_ = 13.136), rFFA (F_4,36_ = 35.567), lOFA (F_4,36_ = 7.56), rOFA (F_4,36_ = 22.866), lSTS (F_4,36_ = 7.98), rSTS (F_4,36_ = 5.637), the rAIT (F_4,36_ = 4.174). For these ROIs, paired sample post-hoc t-test (p<.05, corrected) showed that the 40% phase coherence is always significantly larger than 0%.

There was a significant (p <0.05) task by condition interaction term in the lFFA (F_4,36_ = 4.96), rFFA (F_4,36_ = 3.311), the rSTS (F_4,36_ = 3.620) and the rAIT (F_4,36_ = 3.84), indicating that amplitude increase during detection relatively to fixation was different for different phase coherence levels. Post-hoc t-tests carried out across tasks, within each condition revealed that for these ROIs only the 20% phase coherence conditions were always significantly (p<.05 corrected) larger than all other conditions. The t-values (lFFA (t(9) = 4.407), rFFA (t(9) = 4.132), rSTS (t(9) = 4.684) and the rAIT (t(9)=3.376)) and related effect sizes (Hedges g*: lFFA: 1.361; rFFA: 1.795; rSTS: 1.791; and rAIT: 1.151) further indicate reliable and replicable effects with the current N = 10.

#### Functional Connectivity Measures

Functional connectivity significantly increased (p <0.05, corrected) between multiple areas during the face detection task relative to fixation in multiple core face processing areas. Specifically, we found significant increases in functional connectivity between the 1) lFFA and the rFFA, right and lOFA, rSTS and rAIT, 2) rFFA and rOFA, rSTS and rAIT, 3) the lOFA and the rSTS 4) the rOFA and the rSTS (symmetrical connectivity between regions not repeated).

Multidimensional scaling (MDS), which is a dimensionality reduction technique that allows visualizing the level of similarity amongst data points, was used to summarize task demands related modulations in connectivity. MDS spatial arrangement, where proximity amongst point indicates similarity of responses, highlights how the rAIT is more closely located to the FFAs (i.e. significantly more correlated) during face detection relative to fixation (see Figure 4 Bottom; multidimensional space of fixation rotated onto that of detection using Procrustes transformation).

#### Brain-Behavior Correlations

Figure 5A shows the correlation between the ambiguity score and the difference in the measured brain activity between the stimulus-relevant (SR) and stimulus-irrelevant (SI) tasks. Correlations where significant (p<0.05, corrected) in the bilateral FFA and OFAs and the rAIT.

**Figure 5.**
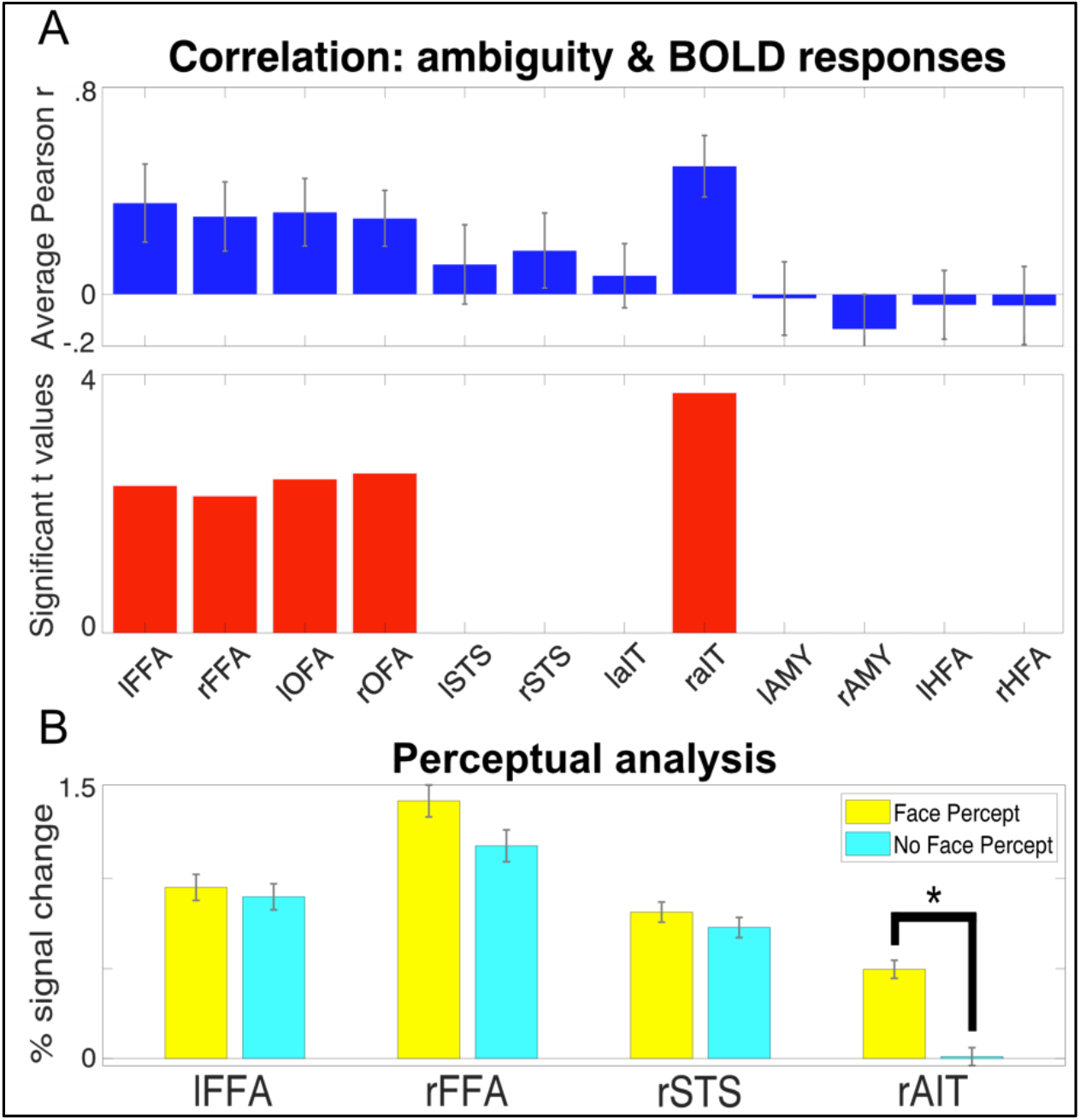
**A)** Brain-Behavior Correlations in FFAs, OFAs and rAIT. **Top**. The Pearson correlation coefficient between the ambiguity score and the difference in task BOLD responses. **Bottom**. For the regions reaching significance (p<0.05), the left and right FFAs and OFAs, and rAIT, the t values are plotted. **B)** Average BOLD response to 20% phase coherence (i.e. the most ambiguous stimulus) for the trials in which participants reported a face percept (yellow) and no face percept (magenta). Responses are reported for the regions that showed a significant (p<.05 corrected) task by condition interaction. Star symbol indicates significantly (p<.05 corrected) difference in amplitude.

#### Perceptual analysis

For the 20% phase coherence condition only (i.e. the most ambiguous percept) we further separated the activity elicited by each trial according to participants perceptual response, creating 2 new conditions: “face percept” and “no face percept”. We investigated amplitude differences between these 2 conditions for those ROIs that showed a significant task x condition interaction. While all areas showed slightly larger BOLD amplitude for the face compared to the non-face percept, paired sample t-tests indicated significant amplitude differences as a function of percept in the rAIT only (t(9) = 6.365 p<.05 corrected; see figure 5B). Moreover, paired sample t-tests contrasting the activation of each condition for all ROIs against 0 further indicated significant (p<.05 corrected) above baseline activation for all ROIs and conditions, except for the “no face percept” response in the rAIT, which did not significantly differ from baseline.

## Discussion

### Task Modulations & Ambiguity

Straightforward manipulation of task demands can produce changes in the amplitude of BOLD response in higher level areas, such as the FFA (Druzgal and D’Esposito, 2001; Kay and Yeatman, 2017; Vuilleumier et al., 2001; Wojciulik et al., 1998) or STS (Narumoto et al., 2001). Traditionally, however, these tasks manipulations are often indirectly related to the neural processing of the stimuli at hand, as they serve a more general purpose of directing attention towards (e.g. N-back) or away from the stimuli (e.g. fixation tasks, see Bokde et al., 2005; Druzgal and D’Esposito, 2001; Egner and Hirsch, 2005; Kay and Yeatman, 2017; Wojciulik et al., 1998). Changes in neural responses to identical visual inputs related to these unspecific changes in task demands often reveal broad contributions from attentional networks, including frontal and parietal regions (Szczepanski et al., 2013). While these manipulations can shed light upon the neural basis of general, non-specific top-down mechanisms, such as awareness, working memory demands or vigilance, they may fail to reveal fine-grained top-down modulations pertaining to the processing of a specific stimulus category. The approach used here instead builds on task manipulations that tap into a relevant stimulus dimension. This disambiguates the contributions of various regions *within* dedicated networks (here, the face processing network), by modulating the difficulty within a stimulus relevant task. In the context of this work, we will be referring to the modulatory forces that direct attention towards (e.g. N-Back) or away from (e.g. observe fixation) the stimuli as *“non-specific top-down”;* and to the task-specific modulations, as they pertain to task difficulty or cognitive load, as *“domain-specific top-down”*.

#### Task modulations: differentiating domain-specific vs non-specific top-down

Our finding that the specificity of task-related modulations is dependent on the nature of the task is consistent with a large body of literature highlighting the conceptual and anatomical differences between different types of attention. Posner and Peterson (1990) delineated 3 attention subsystems devoted to orienting, detection, and alertness. *Our results suggest that a design in which domain-specificity difficulty is varied is capable of revealing the distinct modulation associated with different attentional subsystems*. The significant (p<.05 corrected) main effect of task, indicates, in accordance with previous reports (Druzgal and D’Esposito, 2001; Kay and Yeatman, 2017; Vuilleumier et al., 2001; Wojciulik et al., 1998) that BOLD amplitude is on average larger during the stimulus relevant (SR) relative to stimulus irrelevant (SI) task in canonical face processing areas: the bilateral FFA, OFA and the rSTS. In line with previous work, (Druzgal and D’Esposito, 2001; Kay and Yeatman, 2017; Ress et al., 2000), we suggest that these increases reflect broad top-down contributions relating to increased vigilance and awareness of the stimuli, which have previously been shown to improve decoding performance (Dobs et al., 2018).

By contrast, there are a specific subset of areas that show an interaction between task and stimulus: the bilateral FFA as well as the rAIT and rSTS. Of these only the rAIT showed a significant interaction, driven by the activity elicited by the 20% phase coherence images being larger during the SR compared to the SI. This result indicates that, at least within the context of this work, this region’s responses are *exclusively domain-specific top-down modulations*. In this region in fact, other than for the 20% phase coherence, task demands do not alter BOLD amplitude to any other condition (Figure 3B).

Finally, a large number of regions, specifically the FFAs, OFAs, STS, and rAIT showed a main effect of condition, driven by larger amplitude for the 40% phase relative to the 0% phase. As this comparison is equivalent to a classical ‘face vs scrambled face’ linear contrast (Chen et al., 2007; Rossion et al., 2012) used to define stimulus selectivity, and consistent with the preferentiality of these regions for this stimulus category, this result is unsurprising and won’t be discussed further.

#### Ambiguity

Post-hoc t-tests carried out on the significant task x condition interactions revealed that in all cortical regions exhibiting specific top-down modulations, the stimulus relevant task amplitude increases were always significant *only* for 20% phase coherence. Behaviorally, this condition was also found to be the most perceptually ambiguous, i.e. the condition with the largest reported number of contrasting percepts (i.e. “face” and “no face”). In the context of this study we mathematically defined ambiguity as the inverse of the absolute difference from the inflection point of the sigmoid that describes the face percept/detection behavioral responses (see Equation 1). Both extremes of the phase coherence continuum therefore represent non-ambiguous percepts. That is, at low phase coherence (e.g. 0%) participants consistently reported no face percept and at high coherence (40%) participants consistently reported the presence of a face (Figure 3A). At the maximally ambiguous, 20% condition, the percept was at its most unstable and thus most difficult to categorize, leading to a larger amplitude response. This relative increase is consistent with prior work showing that task difficulty can modulate the BOLD response not only in frontal or parietal regions (Culham et al., 2001; Gould et al., 2003) but also in these category sensitive visual areas (Druzgal and D’Esposito, 2001).

Notably, the reaction times measured in this study do not appear to capture this difficulty increase (Figure 3A). This disconnect between reaction time and experimental performance has been noted before in relation to accuracy and attentional cueing (van Ede et al., 2012) or to task difficulty across a wide range of N-back conditions (Lamichhane et al., 2020). In our data, this likely reflects a disconnect between the idea of “task difficulty” and “difficulty in categorization” which may provide separate contributions to total reaction time.

#### Top-down driven network reconfiguration

To understand how top-down/context modulated the inter-regional neural communication and thus the connectivity within specialized cortical networks (here, the face network), we compared functional connectivity between tasks (see paragraph 4.6.2). By computing functional connectivity on concatenated FIR curves, we: 1) discard the contribution of ongoing activity (which is not the focus of this specific work); and 2) increase statistical power and thus the reliability of our correlational metric. Broadly speaking, connectivity was significantly greater in core and extended regions (i.e. between FFAs, OFAs and rSTS) during the stimulus relevant task, indicating greater communication among these areas, presumably to fulfill task demands. In particular, we observed significantly (p<.05 corrected) greater connectivity between the rAIT and the FFAs during the face task relative to the fixation (Figure 4). A number of studies have suggested a functional differentiation between the right and lFFA (Meng et al., 2012; Rossion et al., 2000), with the former being more involced in holistic processing and the latter in featural processing. It is therefore likely that the degraded stimuli used here (Figure 1) drive both individual feature detection as well as holistic face processing; or that subjects flexibly adapt their strategy depending on the available information, switching from featural (e.g. looking for eyes) to holistic detection, thus engaging both FFAs.

Within the current experimental settings, our data suggest that, during face processing, the rAIT can be recruited by the cortical face network to add an additional resource to resolve ambiguous percepts. These results are consistent with the functional differentiation between core and extended cortical face networks, according to which the core system mediates the representation of more basic aspects of face processing, while the extended system is involved in higher level cognitive functions and can be selectively recruited to act in concert with the regions in the core system (Haxby et al., 2000).

Moreover, our results suggest that network connectivity is not static, but flexibly adapts to the contextual demands. This is in line with an ever growing body of literature advocating the plastic nature of functional connectivity during task and at rest (Allen et al., 2014; Cribben et al., 2013; Debener et al., 2006; Doucet et al., 2012; Hutchison et al., 2013; Sadaghiani et al., 2009) in response to cognitive and behavioral demands (Gratton et al., 2018; Hutchison and Morton, 2016). Our results further expand these views, indicating that even subtle changes in task demands, as those implemented here, can have a dramatic impact over local network reconfigurations.

### Using Stimulus Ambiguity to Differentiate Functional Architecture

Though it is challenging to establish causality, taken together, our results suggest that the rAIT is the source of top-down modulation. First, the correlation between the difference of the BOLD signal between tasks, and each participant’s behavior (ambiguity score) was only significant in the bilateral FFAs and OFAs and the rAIT. By using task differences in BOLD responses to identical stimuli, we sought to highlight task specific top-down effects. This finding, indicating correspondence between the ambiguity function and the difference in BOLD amplitude across tasks within these ROIs, suggests that this ambiguity signal originates from one of these regions. As the rAIT is further along the information hierarchy, it is the plausible source of this signal. Moreover, functional connectivity analysis shows increased connectivity between the rAIT and both FFAs, between OFAs and FFAs, but not between the rAIT and either of the OFAs. In addition, the FFAs show both non-specific (significant main effect of task) and domain-specific (significant task x condition interaction) top-down effects, while the rAIT shows only the task-specific, ambiguity related, top-down effects (significant interaction).

Though prior work has often implicated frontal regions or parietal areas as being a source for these top-down signals when detecting objects such as faces or houses (Baldauf and Desimone, 2014; Kay and Yeatman, 2017), these studies used indirect methods (i.e. 1-back; house or face) to examine face detection. By manipulating difficulty within the context of face detection, we uncover a different, within-network source of top-down modulation by directly stressing the face processing system. This approach is similar to prior work finding modulation within the ventral temporal cortex when viewing degraded facial stimuli (Fan et al., 2020).

Furthermore, there is evidence that the AIT is involved in resolving difficult stimuli in both macaques and humans. For the former, prior work has implicated this region in learning ambiguous stimulus rules related to concurrent discrimination (Bussey et al., 2002). Increases in ambiguity were associated with worse performance in the macaques with perirhinal cortex (i.e. anterior inferior temporal lobe) lesions. In humans, there is evidence that the AIT is associated with discriminating individual face identities (Nasr and Tootell, 2012; Zhang et al., 2016), however these studies employed tasks that did not necessarily tap directly into stimulus-specific dimensions. Though these studies offer support for our findings, they are unable to disambiguate the relationship between behavior and perception, as they either used a memory task, in which subjects could have used an image matching strategy (Nasr and Tootell, 2012), or used only a fixation task (Zhang et al., 2016). Additional evidence for the involvement of the AIT in ambiguous or difficult stimuli comes from outside the face perception literature: this area is active when integrating conceptual and perceptual information (Martin et al., 2018) and it is associated with identifying confusable objects (Clarke and Tyler, 2014; Tyler et al., 2013). We therefore argue that the results presented here and those reported in previous work represent indication that the ambiguity-related top-down signal during face detection originates in the rAIT, is fed back to the FFAs and, from there, to the OFAs. Future work can explicitly assess this using specifically tailored research methods that provide better evidence of causality, such as depth dependent fMRI analyses (Huber et al., 2017), intracranial EEG or fMRI with dynamic causal modeling (Friston et al., 2019).

#### Neural Correlate of Perception

While FFAs and rSTS show both a main effect of task and a task by condition interaction, only the latter was significant in the rAIT. To better understand these differences and further characterize the response profile of these regions, we grouped BOLD responses according to each participant’s percept during the most ambiguous condition (i.e. 20% phase coherence). This analysis shows that, *for the rAIT only*, responses elicited by face percepts are significantly larger than those elicited by no face percepts (Figure 5B), indicating a high correspondence between neural and behavioral responses. Moreover, unlike FFAs and rSTS, showing significant above baseline activation for the no face percept condition, the rAIT shows no significant activation when participants failed to perceive a face (Figure 5B). Taken together, these findings represent evidence for a direct link between the rAIT and subjective perception. These observations are consistent with the idea that anterior temporal regions are involved in subjective awareness (Li et al., 2014) and with reports that the AIT organizes visual objects according to their semantic interpretations (Price et al., 2017). These findings are also consistent with the data from the face localizer (Figure 1) in which the rAIT shows the most preferential face response amongst these 4 regions, with objects and scrambled images not being significantly different from zero. These effects were not found for the lAIT, suggesting a possible lateralization of these processes in humans. However, in light of the low and inconsistent responses in lAIT, we advocate caution in interpreting these results. Further work is therefore required to characterize the functional role of this and other areas (i.e. HFAs and amygdalae) that displayed comparable responses.

The significant activation for the no face percept condition in the FFAs and rSTS can be related to the bottom-up physical properties of the stimulus: regardless of the perceptual state, the 20% phase coherence images always contained a face stimulus. Alternatively, this result can be due to the contextual top-down induced by task demands (here a face detection task). That is, as tasks were blocked by run, during SR runs, participants expected having to resolve perceptual judgment of an ambiguous stimulus, and therefore looked for a face or a face feature in every trial. This is consistent with the observation that, in the absence of a face, the FFA can be activated from contextual cues alone (Cox et al., 2004).

Past work using ambiguous facial stimuli in the form of Mooney faces (Mooney, 1957) have found amplitude differences in the FFA (Andrews and Schluppeck, 2004) or latency differences in the rOFA (Fan et al., 2020). Alternative approaches using bistable perception, such as the face/vase illusion also report that the FFA shows greater activation when faces are perceived (Andrews et al., 2002; Hasson et al., 2001). With binocular rivalry, in which alternative images are shown to each eye, the FFA also increases in activity when faces are perceived (Tong et al., 1998). In our present work we found that the FFA did show a larger, however, non-significant increase when subjects reported seeing a face. This apparent discrepancy with prior reports can be explained by differences in experimental paradigms. Unlike Mooney faces, or face-related bistable stimuli, the 20% phase coherence stimuli used here always have a physical face present. For binocular rivalry, the perceptual ambiguity is similar to bistable perception, however binocular rivalry is ecologically implausible. The defining characteristic of these probes is a dynamic switching amongst percepts despite identical visual input. Here, we have instead focused on the ambiguity of the initial percept, which is more similar to approaches using degraded or partially occluded static stimuli (Flounders et al., 2019; Frühholz et al., 2011). With our stimuli and experimental manipulations, *we isolate a unique and distinctive signal only in the rAIT that distinguishes the subjective perception of a face*.

Another important implication of the findings presented is their interpretation in relation to non-human primate research. Similar to humans, macaque monkeys possess a specialized cortical network in inferotemporal cortex dedicated to processing faces (e.g. Tsao et al., 2003), however the exact correspondence between human and macaque face-selective areas is still unclear (Tsao et al., 2008a, 2006, 2003). While a degree of structural and functional correspondence has been achieved with regards to the core regions (e.g. macaque middle face patches to human FFA (Rajimehr et al., 2009; Tsao et al., 2008a)), identifying a human homologue of the macaque anterior medial (AM) patch (the anterior most patch) has been challenging. Tsao, Moeller, et al. (2008a) for example, failed to uncover a comparable region in humans, attributing such failure to suceptibility related signal drop out due the putative proximity of this area to the ear canal. Rajimehr and colleagues (Rajimehr et al., 2009) uncovered a face specific region in human AIT in 5 out of 10 participants, but were unable to elucidate the nature of its response properties. Here not only were we able to identify a human face selective region in AIT in all subjects, but, importantly, we were able to define its response profile. Our results, linking the function of the face AIT to subjective perception, are in line with recent electrophysiology reports, showing that activity in the AM face patch is related to the animal’s individual ability to detect a face (Moeller et al., 2017).

### Benefits of High Field Imaging

As briefly mentioned in the above paragraph, much of the difficulty in imaging regions such as the AIT relates to low SNR associated because of susceptibility artifacts and inefficient RF transmit fields in the ventral temporal lobes (Devlin et al., 2000). Moreover, areas such as the AIT are typically difficult to image at more conventional (3T) field strengths due to low signal and are often not located in every individual (e.g. 50% in Rajimehr et al., 2009), and often show large deviation from expected hemodynamic responses (e.g. Ramon et al., 2015). These issues can persist independent of field strength.

Here we used UHF fMRI to capitalize on higher SNR, CNR, and acceleration performance maximizing fMRI sensitivity in these regions. It should be noted that moving to higher field alone is not sufficient to guarantee increased image quality in these regions. At UHF, while BOLD signal changes and SNR do increase, B0 and B1+ inhomogeneity also increase and have to be dealt with. Here the combination of our experimental design, rigorous analytical approach, manual B0 shimming tailored to all participants with a specific focus on anterior ventral temporal regions and flip angle optimizations (see methods) yielded fruitful results. For all our participants we report consistent and corresponding regions in the anterior inferior temporal cortex on both the left and right that preferentially respond to faces and that are modulated by task demands. This increased sensitivity that led to large effect size and ROI identification in all subjects, stems from a combination of parameters and sequence optimization. Specifically, in addition to the increased SNR that accompanies UHF strength, unlike 3T acquisitions, where no effort has been put forward to for optimizing B0 and flip angles, we manually adjusted B0 inhomogeneity and flip angles to maximize SNR for each subject. Moreover, relative to previous human work, where functional voxels measured > 3 mm iso (e.g. (Rajimehr et al., 2009; Ramon et al., 2015)), here, we used 1.6 mm iso voxels, minimizing partial volume effects and spin dephasing, ultimately reducing signal loss in dropout regions (Thanh Vu et al., 2017). While the increased signal due to higher field strengths can be a benefit, using high field alone is insufficient and can be, in fact, at times detrimental. Appropriate consideration of tradeoffs, such as increased B0/B1^+^ instability, is necessary.

### Beyond the ventral temporal lobe: future directions

Beyond the ventral temporal lobe, there have been reports of face sensitive regions in lateral orbital regions in macaques (Barat et al., 2018; Hadj-Bouziane et al., 2008; Tsao et al., 2008b). However, evidence for the existence of these areas in humans has been mixed, uncovering such regions in approximately half (Troiani et al., 2016) or one-third of the study population (Troiani et al., 2019; Tsao et al., 2008a). In the current work, we are able to identify these areas in all subjects during the localizer portion of the study. Activation was proximal to the H-Shaped sulci, consistent with both prior reports (Troiani et al., 2019) and across subjects, as visualized after normalization to MNI space (Figure 2). Like the other ROIs in the face preferential cortical network, these regions exhibited positive responses to faces and significantly (p< .05) greater activity to faces relative to scrambled stimuli or objects (See Methods).

Although we did not observe meaningful stimulus relevant task modulations we did observe an interesting, although non-significant, separation of 2 areas, with regard to their functional connectivity during face detection: the rHFA exhibited a non-significant increase in activation, while its left counterpart exhibited a non-significant decrease. These results should be interpreted with caution, as the primary tasks produced low response amplitudes relative to the localizer. The absence of strong activation in the HFAs, as well as other regions such as the amygdalae and left AIT, during the primary task despite prominent responses in the localizer task is likely due to a number of differences between the tasks. These include, but are not limited to, difference in presentation timings (i.e. 12 secs vs. 2 secs on/off) and the fact that the localizer used stimulus presentations with non-degraded, full faces with variable expressions and gaze directions.

That these task differences drove larger responses during the localizer task in the HFAs is congruent with prior findings, namely that these areas have been suggested to be involved in social and emotional aspects of face processing in non-human primates (Barat et al., 2018), aspects that, within the primary task context, are irrelevant. Our failure to find task modulations is therefore consistent with the more complex evaluative role of the frontal cortices (Noonan et al., 2012) and its engagement in social perception tasks (Barat et al., 2018; Beer et al., 2006; Freeman et al., 2010; Mah et al., 2004).

Given the enlarged frontal cortices among primate species and in particular the highly developed frontal areas in humans, it will be essential to perform further research that build on these differences and further manipulate context, value and/or salience in order to elucidate the functional role of regions such as the HFAs during face perception and processing.

## Conclusion

Using faces, a well-studied stimulus underpinned by a dedicated, complex cortical network, in this study we combined high SNR fMRI with domain-specific attention and observed 2 types of top-down scaling mechanisms: 1) a broad gain effect related to drawing attention to the stimuli, manifesting as an unspecific amplitude increase across conditions; and 2) an additional scaling of specific conditions dictated by domain-specific task requirements. We explain the latter in terms of perceptual ambiguity and suggest that the ambiguity signal originates in the rAIT. Importantly, only in the rAIT is both preferentially active under challenging conditions and predictive of the subjects’ perceptual judgments. We further show that subtle changes in task demands can lead to dramatic changes in network reconfigurations. Our results suggest that the combination of an explicit face detection and stimulus matched control task with low noise fMRI capable of resolving previously inaccessible regions of the human brain may be the only way to understand the changes underlying human cognitive flexibility

## Acknowledgments

Funding for this study was provided by National Institutes of Health Grants RF1 MH117015 (Ghose), RF1 MH116978 (Yacoub), and P30 NS076408 (Ugurbil). The authors would also like to thank Dr. Kendrick Kay for conceptual discussions regarding stimulus ambiguity and feedback.

